# Co-occurrence is associated with horizontal gene transfer across marine bacteria independent of phylogeny

**DOI:** 10.1101/2025.03.25.645238

**Authors:** Gavin M. Douglas, Nicolas Tromas, Marinna Gaudin, Patrick Lypaczewski, Louis-Marie Bobay, B. Jesse Shapiro, Samuel Chaffron

## Abstract

Understanding the drivers and consequences of horizontal gene transfer (HGT) is a key goal of microbial evolution research. Although co-occurring taxa have long been appreciated to undergo HGT more often, this association is confounded with other factors, most notably their phylogenetic relatedness. To disentangle these factors, we analyzed 15,339 marine prokaryotic genomes (mainly bacteria) and their distribution in the global ocean. We identified HGT events across these genomes and enrichments for functions previously shown to be prone to HGT. By mapping metagenomic reads from 1,862 ocean samples to these genomes, we also identified co-occurrence patterns and environmental associations. Although we observed an expected negative association between HGT rates and phylogenetic distance, we only detected an association between co-occurrence and phylogenetic distance for closely related taxa. This observation refines the previously reported trend to closely related taxa, rather than a consistent pattern across all taxonomic levels, at least here within marine environments. In addition, we identified a significant association between co-occurrence and HGT, which remains even after controlling for phylogenetic distance and measured environmental variables. In a subset of samples with extended environmental data, we identified higher HGT levels associated with particle-attached bacteria and associations of varying directions with specific environmental variables, such as chlorophyll *a* and photosynthetically available radiation. Overall, our findings demonstrate the significant influence of ecological associations in shaping marine bacterial evolution through HGT.

## Introduction

Closely related prokaryotic taxa often display substantial differences in gene content, largely due to horizontal gene transfer (HGT) [1]. To better understand these dynamics, gene transfer networks have been investigated across diverse species [2]. A recurrent observation from this work is that taxa from similar environments are associated with higher rates of HGT [3–6].

This observation is expected, for at least two reasons. First, all HGT mechanisms are dependent on physical proximity to some degree [7]. For instance, conjugation requires a physical interaction between cells for DNA transfer to occur. Second, taxa from the same environment can experience similar selective pressures and may potentially benefit from the same set of adaptive genes. This is clear within species, where strains experiencing similar pressures exhibit parallel adaptive point mutations [8]. It is expected that similar dynamics occur between strains within the same environment based on horizontally-transferred genes [9]. Genes can also spread horizontally between more distantly related taxa within a community. This is clearly seen for antibiotic and heavy metal resistance genes, which have the potential to confer adaptive benefits to many taxa in a contaminated environment [10].

Although these explanations for the association between HGT and co-occurrence are intuitive, there is also an important confounder to consider: phylogenetic distance. This point was recently raised in work disentangling the action of these factors in driving HGT inferred across reference genomes from isolates, and based on co-occurrence patterns of 16S rRNA gene sequences [11]. In this work, the authors found that the positive association between co-occurrence and horizontal gene transfer remained after controlling for phylogenetic distance, although the association was decreased.

This study is an important contribution but was limited to 16S rRNA gene data used for identifying co-occurring taxa. As this is a coarse approach, it cannot reliably identify strains or species within a sample. In contrast, shotgun metagenomics provides improved resolution for identifying strains, given sufficient genome coverage [12]. In addition, due to the broad and cross-environment focus of this study [11], the authors did not thoroughly investigate how particular environmental factors are associated with HGT.

Marine microbial communities are an appealing system for studying prokaryotic evolutionary dynamics. Marine prokaryotes undergo widespread HGT and are stratified into communities based on depth and other environmental factors [13–15]. In addition, understanding prokaryotic community dynamics and evolution is important as marine microbes play key roles in global biogeochemical cycles [16], and are also experiencing heightened environmental stress due to climate change [17]. To help better understand these ecological and evolutionary dynamics, numerous ocean samples have undergone metagenomics sequencing [14, 18].

Here, we used publicly available ocean metagenomes to investigate patterns of HGT and co-occurrence across the global ocean. Our key question focused on whether the link between HGT and co-occurrence in this dataset is robust to the inclusion of phylogenetic distance as a covariate. While this question was also recently investigated across biomes [11], here we specifically focus on marine samples with shotgun metagenomics data. Another novel aspect of our work is the consideration of additional factors that could be driving the association between HGT and co-occurrence. In particular, by using environmental data associated with these metagenomics samples, we investigated to what degree shared environmental pressures can explain the observed association between HGT and co-occurrence. We also consider a wide range of methods for inferring HGT and co-occurrence to ensure our results are robust.

## Methods

### Acquiring genomes

We downloaded all 34,815 genomes from a prior metagenomic study of the global ocean microbiome [19]. This included whole genome sequencing from isolates, single-amplified genomes, and metagenome-assembled genomes (MAGs), along with pre-existing Prokka gene annotations [20], taxonomic classification, and quality reports.

These data correspond to version 1.0 of the Ocean Microbiomics Database, which we refer to as our focal dataset. We used the existing taxonomic classifications [19], which were previously obtained using the Genome Taxonomy Database Toolkit (GTDB-Tk) [21]. Using quality metrics provided with these data, we retained 15,339 genomes with mean completeness > 75% and mean contamination < 5%. The contamination cut-off was chosen based on the established cut-off used for defining high-quality draft genomes [22]. False positive HGT events would be driven primarily by contaminated, rather than incomplete, genomes, so we used a more relaxed completeness cut-off of > 75% (rather than the typical > 90% for high quality genomes [22]) to include more genomes in our analyses. This final set corresponded to 11,833 MAGs, 1,862 single-amplified genomes, and 1,644 genomes from whole genome sequencing of isolates.

### Genome phylogenies

GToTree v1.7.00 [23], with default parameters, was run on the 15,339 genomes using the Bacteria and Archaea Hidden Markov Model-set composed of 25 single copy genes to generate an alignment protein file with HMMER v3.3.2 [24, 25] and Muscle v5.1 [26]. Automated trimming was performed on the alignments using trimAl v1.4 [27], with the - automated1 and -phylip options. IQTREE2 v2.0.7 [28] was then used to construct a phylogenetic tree, using ModelFinder Plus and 1,000 bootstraps. The best-fitting model, ‘Q.yeast+I+R10’, was automatically chosen based on a comparison of Bayesian Information Criterion values. Pairwise (patristic) distance between tips was computed using the castor R package v1.8.2 [29].

To compare our focal results (heavily based on MAGs) to those based on a set of higher-quality genomes, we also downloaded all representative aquatic genomes from the proGenomes database v3 [30]. Taxonomy for each genome was parsed based on TaxIDs using the taxize R package v0.9.102 [31]. Each genome in proGenomes is associated with a BioSample in the National Center for Biotechnology Information database, which we used to parse sample metadata using efetch v21.6 [32]. All genomes linked to metagenomic samples or without any contigs greater than 10,000 bp were excluded, to help avoid misassembled genomes. This filtering resulted in a final set of 5,551 genomes (out of 11,060) for this database. The same workflow as above was used to produce a phylogenetic tree, based on the same best-fitting model, and pairwise distances.

### Metagenomic sample preprocessing

To characterize the distribution of our filtered genomes across environments, we also prepared a metagenomics dataset representing diverse ocean samples that were then mapped to these genomes. To that end, we acquired all 1,865 ocean water metagenomics samples listed in the OceanDNA resource [33]. This raw metagenomics sequencing data was downloaded from the National Center for Biotechnology Information with Globus Connect [34]. We filtered out low-quality reads with fastp v0.23.4 [35], with the options -l 70, --cut_mean_quality 20, --cut_front, --cut_tail, --cut_window_size=4, and --detect_adapter_for_pe (this last option only for paired-end FASTQ files). Files corresponding to technical replicates were concatenated. We then mapped all filtered metagenomics reads to the genomes with CoverM v0.6.1 [36], using the options --exclude-supplementary, --min-covered-fraction 0, and --proper-pairs-only (for paired-end reads). We called genomes as present based on a >= 30% breadth of coverage cut-off (i.e., where at least 30% of a genome was mapped by at least one read). For each sample, we set the reads per kilobase per million mapped reads values for genomes with breadth below the presence cut-off to 0. We excluded genomes not called as present in at least 10 samples, which resulted in 10,388 remaining genomes. We excluded three metagenomics samples where no genomes were called as present.

To distinguish metagenomics samples more likely to represent free-floating microbes, compared to particle-attached microbes, we partitioned samples based on the size-filtration steps used during sample collection, which are commonly referred to as size fractions. A cut-off of 3 µm is commonly applied to distinguish free-floating (small particles) and particle-attached (large particle-associated) microbes by size [37–39].

This approach is a proxy for habitat preferences, which could correspond to more planktonic (free-floating) or particle-associating lifestyles, but this filtering process is imperfect and should not be taken as absolute classification. We categorized small particle samples as those size-selected at <= 3 µm, which corresponded to 906 samples. However, as this was not the initial focus of our analysis, we did not acquire metagenomics data equally split by size-selection criteria, and almost all samples did not exclude small particles. Thus, it is not possible to categorize our initially analyzed metagenomics samples as primarily particle-attached. Instead, we opted to categorize 846 samples as “less-filtered”, which are those with an upper size-selection filter >= 20 µm. Our rationale for this category is that they are more likely to contain DNA from particle-attached prokaryotes compared to samples filtered to enrich for small particles. We then identified cases where genomes were predominately found in either small particle fraction size or “less-filtered” (large and small particle size) groups. We classified genomes as being associated with a given group if > 75% of the samples containing the genome were in that group. We also required that the median relative abundance of this genome, based on reads per kilobase per million mapped reads, was highest in this sample subgroup. Based on this criterion, 346 focal genomes could be classified as enriched in “less-filtered” and 2,470 as enriched in small particle metagenomics samples, respectively. We applied this same approach for the proGenomes database as well, where we could classify only 24 genomes as “less-filtered” and 137 as small particle-associated.

The OceanDNA resource lists several independent datasets. To ensure our results were not driven by technical differences between these datasets, we also subdivided metagenomics samples to be only those part of the *Tara* Oceans campaign (n=368) or the GEOTRACES campaign [40] samples (n=610).

After our initial analysis we conducted a follow-up with 160 additional ocean metagenomes from the *Tara Oceans* campaign that were processed to have size-selection filters ranging from 3 - 20 µm (referred to as large particle samples). These samples were originally meant to capture the protist-enriched fraction of ocean water [41, 42], but also include the size range for particle-attached bacteria. These metagenomic samples were pre-processed and mapped to the genome database with CoverM using the same steps as above, where only samples with at least 10 genomes called as present were retained. We then subdivided to 43 biological samples for which there was a match between the small (0.22 – 3 µm) and large (3 – 20 µm) particle-associated metagenomics sample. All small particle-associated metagenomics samples were already included in the subset of the OceanDNA dataset described above. We focused on this paired-sample approach (specifically for this analysis) to minimize the impact of environmental and technical confounding factors.

### Clustering genes and identifying reciprocal best hits

Genes in the Ocean Microbiomics Database v1.0 that we analyzed were previously clustered based on sequence identity, with a representative gene indicated per cluster. However, these clusters included genes with high variation in length (e.g., genes ranging from 3 kb to 21 kb in the same cluster). Accordingly, we re-ran gene clustering to produce clusters of genes with high pairwise identity over their entire lengths.

To do so, we first parsed out all protein coding genes sequences from pre-existing GFF files per genome. We then clustered these genes with CD-HIT-EST v4.81 [43], with a sequence identity threshold (-c) of 0.95, a length difference cut-off (-s) of 0.95, alignment coverage settings (-aL and -aS) of 0.95, and with accurate mode (-g 1).

Alignments were then built with MAFFT v7.525 [44], with the options --retree 2 -- maxiterate 2 to speed up processing, as these were generally trivial alignments. We then computed the pairwise hamming distance for all sequences per cluster in Python using the hammingdist package (v1.3.0). We identified gene pairs that were reciprocally the closest matches using this approach and between taxa that were at least in different genera (i.e., that were not in the same genus or species). We used a lower sequence identity cut-off of 95% as this allowed us to potentially detect ancient HGT events, in addition to more recent events, which we assumed would be enriched among matches with sequence identity >= 99%. For all HGT-detection approaches, all genes (or hit regions, in the case of the BLASTn approach, described below) on contigs < 5000 bp were excluded after detecting HGT events.

### Sensitive approach for identifying indirect gene flow with pairwise BLAST

In addition to the clustering approach, we also performed pairwise identity comparisons between all genomes using BLASTn, with BLAST+ v2.12.0 [45]. Only hits >= 500 bp long and at least 95% identical were retained. To clarify, BLASTn matches were generally based on short subsequences in contigs, rather than the entire contigs. Our analysis for these data focused only on genomes within different genera or higher (i.e., above the genus-level as for the cluster-based HGT approach), to reduce the number of false positives due to vertical inheritance rather than HGT. We ran bedtools v2.30.0 [46] to intersect gene annotations with BLASTn-identified matching regions. For cases where multiple hits overlapped (for the same pairwise genome comparison) the highest scoring hit was retained.

BLASTn matches were called as putative HGT events regardless of whether they were the reciprocal best hits in the dataset. Thus, this approach is not meant to identify direct HGT events between genomes, and instead identifies regions between genomes that were putatively horizontally acquired through a shared HGT network. We used this information for complementary purposes, and expected these calls to be more sensitive, but also enriched for more false positives, and less straightforward to interpret than the clustering method described above.

### Phylogenetic reconciliation within species

We also identified putative gene transfer events within species with RANGER-DTL [47, 48]. We included this approach as using sequence identity alone is unreliable for detecting HGT between closely related genomes (due to high identity stemming from a recent common ancestor). We also ran HoMer v1.0 [49] in conjunction with RANGER-DTL to call multi-gene transfer events.

We ran this approach over 190 species that had at least 10 genomes. We found that topological differences between the core genome tree and accessory gene trees were widespread: a mean of 91.6% (standard deviation: 0.06%) of tested genes were inferred to have at least one transfer event across all species. We had little confidence in the accuracy of individual events (see Discussion) and instead used these results only for supplementary purposes. Our assumption here is that the relative ranks of inferred and real HGT event counts between genome pairs are highly correlated, even if there is a baseline of noise.

### Functional annotation and enrichment

Clusters of Orthologous Genes (COG) annotations were taken from a previous run of eggNOG [50], which was distributed with the genomes. For each eggNOG Orthologous Group, we parsed out all COG annotations in the definition and then re-mapped them to higher-level COG categories. To determine the COG annotation associated with each gene cluster, we considered the classifications of each gene in the cluster and used the majority rule to determine which COG (and COG categories) to assign. We also annotated all genes that matched to mobile genetic elements in the proMGE v1 database [6]. We first translated all protein coding gene sequences to amino acids with the transeq EMBOSS program v6.6.0.0 [51], using the parameters -table 11, -trim, and - clean. We then ran HMMER v3.1b2 to identify matching genes. These represent a range of transposase and recombinase protein-coding genes associated with diverse MGEs, such as insertion sequences and integrative conjugative elements.

We also ran geNomad v1.5.0 [52] with database v1.1, and default parameters except -- cleanup and --splits8 on the contigs of all studied genomes. This analysis was run independently of the proMGE analysis, and returns MGE-associated contigs rather than specific genes. This allowed us to identify contigs categorized as ‘virus’ or ‘plasmid’, as well as those that contain prophages. We ran all enrichment tests for the MGE categories on genes rather than clusters due to this focus on contig-level annotations.

The expected enrichments of COG categories in the HGT results (prior to cross-referencing with co-occurrence data) were taken from a previous study [11], which itself based these expectations upon results across multiple studies [5, 53–61]. We defined categories expected to be depleted or enriched as those showing the consistent signal in all but one study (excluding studies with no result for a given category).

When testing for functional enrichments among clusters involved with HGT events vs. all other gene clusters, we only considered clusters assigned to a COG annotation. We also excluded rare clusters annotated as eukaryotic-specific COG categories (A, B, Y, and Z). We then ran Fisher’s exact tests on the number of clusters in each COG category in at least one HGT relationship compared to all other clusters in that category, relative to the background of all clusters in other COG categories (significance called at Benjamini–Hochberg-corrected [62] P-values < 0.05). We focused the COG category functional enrichment analysis on clusters, rather than individual genes, so that clusters with many genes involved in HGT events would not be counted multiple times as independent observations.

### Co-occurrence analyses

We used the backbone R package [63] to run hypergeometric distribution-based co-occurrence tests for all pairwise genomes. The exact version 1.3.1 of this package needed to be used, as more recent versions no longer provide the method we required at scale. This allowed us to identify cases where genomes co-occurred more often than expected by chance (i.e., if they were randomly distributed across samples). We computed the effect size as the co-occurrence ratio, the ratio of observed vs. expected number of metagenomics samples containing both genomes. These values were used for comparison with phylogenetic distance, but we primarily focused on the binary classification of significantly co-occurring genome pairs. Significantly co-occurring genomes were identified based on Benjamini-Hochberg corrected P-values < 0.05 and co-occurrence ratios > 1.

To ensure our inferences were robust to this bioinformatic step, we also computed co-occurrence using two other approaches. First, we computed a simple overlap metric, as used previously [11]: 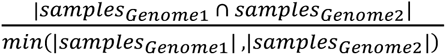. This approach does not adequately correct for the total prevalence of both genomes being compared (unlike the hypergeometric approach above). For instance, if a genome is found across all samples, then it will trivially have a co-occurrence of 1.0 with all other genomes. The second additional approach we used was to compute co-abundance with the propr R package [64] v5.1.4, which incorporates the relative abundance of genomes. This metric was computed using the NetCoMi R package [65] v1.1.0, for convenience.

### Statistical modelling

We ran logistic regression in R using the glm function. These models were based on pairwise comparisons of genomes, with predictors of co-occurrence, tip distance, and the difference in median environmental values of the samples each genome is found in. We also included categorical variables of which filter cut-off grouping the genome pair is within (see Results). For the focal model, the response was the binary value of whether each genome pair was inferred to have at least one HGT event. For models basing co-occurrence upon simple co-occurrence and propr-based co-abundance, these co-occurrence values were continuous rather than binary. All continuous variables were transformed using ordered quantile normalization [66] using the bestNormalize R package v1.9.1. We computed the variance inflation factor for each fixed effect using the performance R package v0.12.4 [67].

As the key data we analyzed was based on genome pairs, we also wanted to control for genome ID, to ensure that a small number of genomes (e.g., in numerous co-occurrence or HGT relationships) were not driving our results. To that end, we also built generalized linear mixed models with the R package glmmTMB v1.1.10 [68], with genome ID as a random effect. For this supplementary analysis, we duplicated the input table for each model, so that each genome comparison was present twice, and assigned once to each genome being compared. Due to the table being duplicated, we do not have confidence in estimates of variability from these models, and used them simply to assess whether our overall interpretations shifted when controlling for genome ID.

### Computing prevalence of genomes with horizontal gene transfer or related features

In addition to our logistic regression models, which focused on genome pairs, we calculated summary metrics that provide insight into the extent of HGT and other related characteristics per metagenomic sample. We did so by identifying all genomes called as present in a metagenome and counting how many had at least one instance of putative HGT called (based on the focal cluster-based approach). We similarly tallied genomes with at least one gene that matched the proMGE database, that were identified as plasmid or virus-related with geNomad, or for which geNomad identified at least one contig as containing a provirus. For each feature, this count was divided by the number of genomes in the sample, to return the proportion of genomes with at least one instance of the feature of interest.

### In-depth environmental data analysis

To explore a wider range of environmental factors, beyond those present in the OceanDNA dataset, we acquired all environmental depth sensor data for *Tara* Oceans samples from the PANGAEA database [69]. For environmental variables with multiple summary statistics, we retained only the median value per sample and based on locally-calibrated estimates where available. We restricted to data intersecting with *Tara* Oceans samples we acquired (e.g., ignored metatranscriptomic data), and then retained only a single representative of highly redundant variables (defined as those with Spearman correlation coefficients >= 0.95). Based on this approach, we excluded: beta470 and bb470 (redundant with bbp470); absolute PAR (redundant with percent PAR); conductivity and potential temperature (redundant with temperature). We also excluded bac660 and bacp due to the difficulty of interpreting these optical properties, which are primarily used as intermediate measures to calculate other properties.

After this pre-processing, we focused on these variables:

- Latitude (degrees)
- Longitude (degrees)
- Depth (metres)
- Temperature (°C)
- Potential density (kg/m³; Sigma-theta)
- Salinity
- Nitrate (NO₃⁻; µmol/L)
- Oxygen (µmol/kg; Calibrated using World Ocean Atlas 2009 climatology)
- Chlorophyll *a* (mg/m^3^; Nonphotochemical fluorescence quenching-corrected and calibrated using water samples). This is a measure of phytoplankton biomass, so it is a biotic variable although we group it with environmental variables herein.
- Photosynthetically available radiation (PAR [%]; Based on diffuse attenuation coefficient and average of eight-day satellite readings)
- fluorescent Coloured Dissolved Organic Matter (fCDOM; Parts per billion quinine sulfate equivalents)
- Backscattering coefficient of particles (per metre) at 470 nm (bbp470). This measure is related to particulate organic carbon levels [70–72]

We focused this analysis on 131 *Tara* Oceans samples that were all from the small particle-associated size fraction, with at least 10 genomes present based on CoverM mapping and with no missing environmental data. We then computed the Spearman correlation coefficients between all environmental variable pairs and computed pairwise distance as |1 – Spearman’s ρ|. We performed hierarchical clustering with the average method and visualized the dendrogram of variable groupings and clustered heatmap of standard scores across all samples with the ComplexHeatmap v2.20.0 R package [73].

We ran a random forest model [74] with 10,000 trees with the ranger R package v0.17.0 [75] to predict HGT prevalence in a metagenome based on these environmental variables. The number of variables to subsample at each split of each tree (the mtry parameter) was set to the default (3, given 12 predictors). Variable importance was computed based on the permutation method. To determine which variables were significantly predictive in the random forest, we re-ran random forests for 1,000 replicates with the dependent variable (HGT prevalence) shuffled. We used the inferred variable importance for each variable across these replicates to build null distributions of the variable importance under no signal. We calculated P-values based on the proportion of simulation replicates that resulted in variable importance equal to or higher than the observed importance for each variable (and called significance at Benjamini-Hochberg-corrected P < 0.05).

We also confirmed our overall variable importance findings were not affected by parameter settings. Specifically, we confirmed that changing the mtry parameter (ranging from 1-11) did not result in substantial changes in the order of variable importance. We also confirmed that including just one of a highly correlated pair of variables (e.g., depth and photosynthetically available radiation) did not result in the other variable’s relative importance substantially changing (i.e., ensured variables were not masking each other). This analysis was conducted post-hoc to confirm model robustness, not for model selection.

We undertook several steps to make our random forest model more interpretable [76], using the iml R package [77] v0.11.4. This included Individual Conditional Expectation plots, which show how the prediction for each sample would change for different values of a predictor (an environmental variable in this case). We also computed pairwise interactions between significant variables based on the H-statistic [78]. This metric represents the proportion of the joint effect of both variables that is explained by their interaction.

### Working environments

Custom code for processing HGT and co-occurrence result tables was written in Python (run with v.3.10). Visualizations and most statistical analyses were conducted in R v4.4.0 [79]. Where possible, Linux commands (on Ubuntu 22.04.5 LTS) were parallelized with GNU Parallel v20240322 [80] and tools were installed from Bioconda [81]. Claude large language models were used to help debug code and to check for errors (in code and manuscript).

### Code and data availability

The accessions of most shotgun metagenomics data used in this study are present in the OceanDNA FigShare repository [33]: https://doi.org/10.6084/m9.figshare.c.5564844.v1. The exception is for the protist/particle-attached fraction samples, which were taken from the BioProjects PRJEB4352 and PRJEB9691. All metagenome-assembled genomes and pre-existing annotations were taken from https://sunagawalab.ethz.ch/share/microbiomics/ocean/suppl_data/. All code and key data files are distributed through a Zenodo repository (https://doi.org/10.5281/zenodo.14867578), which includes the filtered sample metadata and genome information.

## Results

### Identifying putative horizontal gene transfer events

To study prokaryotic genome diversity in the global ocean, we analyzed 15,339 genomes representing 3,270 species, which were previously constructed from marine metagenomics samples [14, 19, 82]. These genomes are from diverse lineages (**Supplementary Figure 1**), primarily bacteria (90.9%). The most common phyla are Proteobacteria (47.9%) and Bacteroidota (15.2%) and the best represented archaeal phylum is Thermoplasmatota (5.9%).

There are several approaches for inferring HGT, which can produce different but complementary results [83]. Accordingly, we identified putative HGT events between these genomes using several methods to ensure the overall results remain unchanged (see Methods). The first two approaches were based on identifying regions with sequence similarity >= 95% between genomes from different genera (or that differ at higher taxonomic levels). In other words, we did not consider genome comparisons within the same genus or species. Our key assumption is that regions of high similarity are unlikely to occur (when of sufficient length) in the genomes of distantly related taxa due to these regions being present in a common ancestor [5]. Instead, we take these observations to represent putative HGT events.

Based on this framework, our focal HGT-detection approach focused on clustering all genes across all genomes (**Supplementary Figure 2**). We identified 9,046 putative HGT events based on this approach, 71.2% of which were >= 99% identical, and for which the majority were between genomes within the same family (**Figure 1a**). Although this is a relatively high number of putative HGT events, the number of events called between each pair of genomes was low (median of 1.0 and mean of 3.95 in genome pairs with at least one putative event).

**Figure 1:**
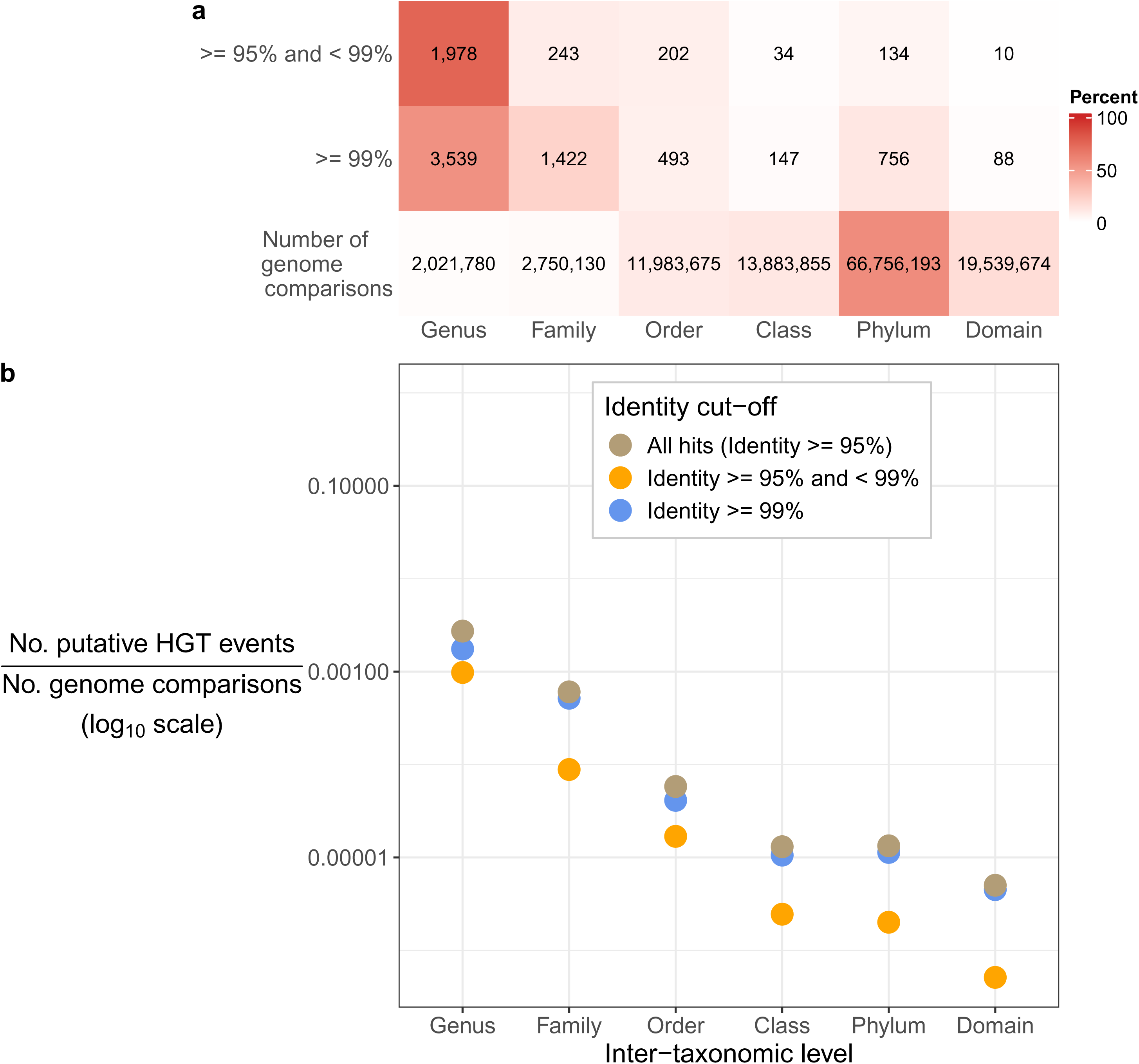
Summary of putative horizontal gene transfer events based on sequence identity cutoffs. (a) Counts of putative horizontal gene transfer (HGT) events based on the cluster-based approach, by the lowest taxonomic level at which the compared genomes differed. The total number of genome comparisons per level is also indicated. These cells are coloured by the percentage per row. (b) Proportion of putative horizontal gene transfer events per taxonomic level, after normalizing by the total number of genome comparisons per level.

We reasoned that the sequence identity of putative HGT hits, on average, would provide information on the age of the transfer event. One expectation of older HGT events is that mutations and recombination will result in a shorter region identifiable as a transfer event. We used a complementary approach based on BLASTn (see Methods) to evaluate how the lengths of matching regions vary by sequence identity. As expected, regions with high sequence identity (>= 99%) were longer and included more genes on average (mean: 6.6; standard deviation [SD]: 6.6) compared to those with lower sequence identity (>= 95% and < 99%; mean: 2.7; SD: 4.2).

We then investigated whether the expected negative relationship between evolutionary divergence and HGT held in our marine dataset. There is a clear signal of decreasing HGT rates at more distant taxonomic levels (**Figure 1b**), after normalizing for the number of pairwise genomes compared at each level. This signal is similar for both the older (< 99%) and more recent (>= 99%) putative HGT events.

### Functional enrichments of horizontally transferred genes

We next asked whether the inferred horizontally transferred genes were enriched in expected functional groups identified in previous studies [11]. Most COG categories were significantly enriched in expected directions (**Figure 2**). For instance, mobile elements (X) and defense mechanisms (V) were strongly enriched among HGT events. In addition, all significant categories that were expected to be depleted for HGT were identified for putative transfers between identities of >= 95% and < 99%. However, transfers at >= 99% identity showed a more mixed signal for categories expected to be depleted for HGT. For instance, we observed a mixed signal of enrichment for lipid transport and metabolism genes (I), and the unexpected enrichment of several other categories, especially at higher taxonomic levels.

**Figure 2:**
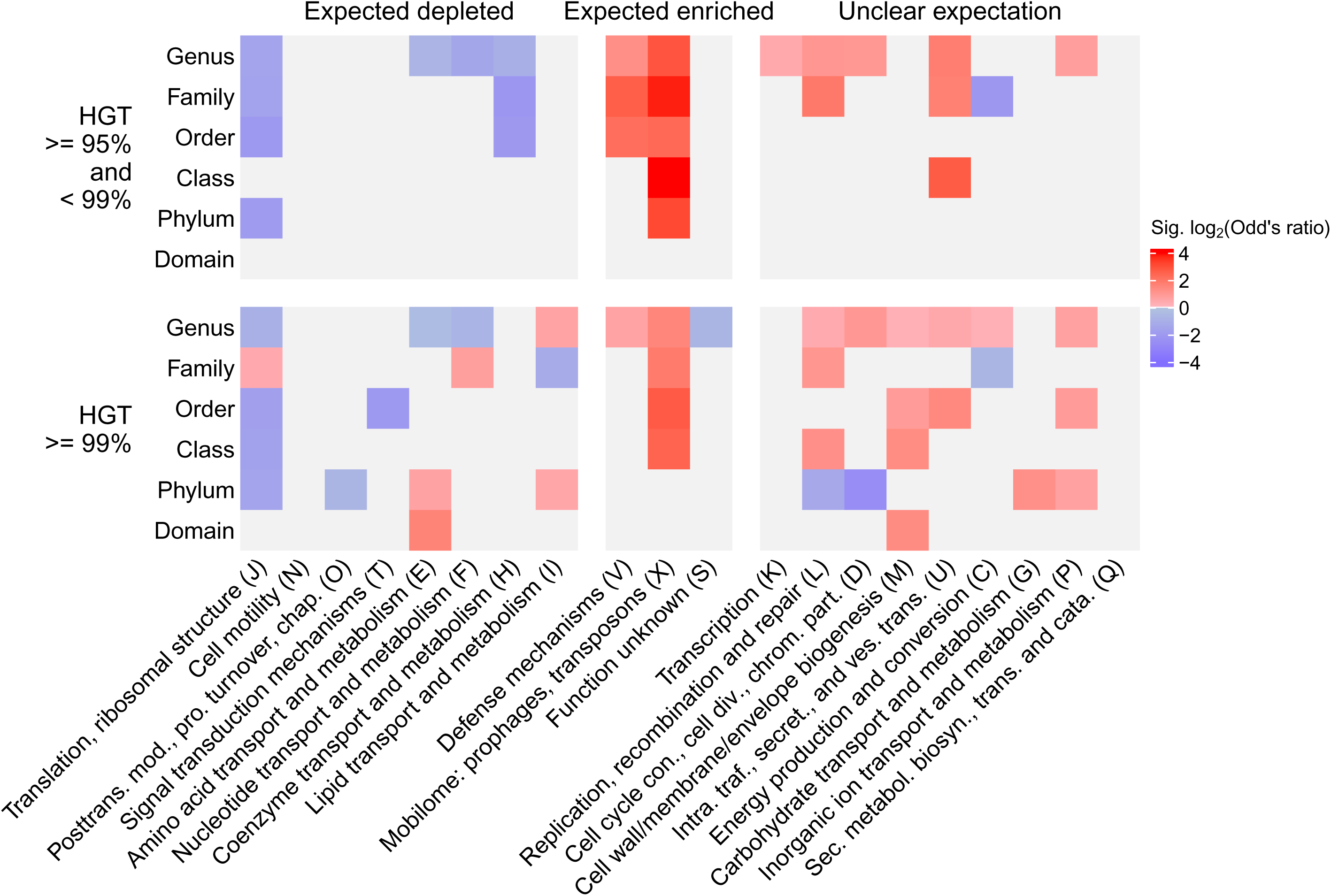
Cluster of Orthologous Genes (COG) categories enriched among horizontal gene transfer (HGT) hits based on sequence identity. HGT results used for this analysis were based on the cluster-based approach. Expected patterns of enrichment are indicated for COG categories at the top, and individual COG categories (in abbreviated form in some cases) are indicated at the bottom. The row groupings indicate the identity cut-off, in addition to the taxonomic level at which the genomes that encode the genes differ. Results are only shown for significant Fisher’s exact tests (Benjamini-Hochberg-corrected P < 0.05); light grey cells indicate non-significant tests.

This observation highlights a unique aspect of our enrichment analyses, which is that we conducted them for inferred HGT events at each taxonomic level separately (**Figure 2**). This allowed us to identify cases where the same category is enriched in different directions at different taxonomic levels, such as those involved in chromosome partitioning and cell division (D) and amino acid transport and metabolism (E).

We also explored mobile genetic element-associated gene annotations and overall found them enriched among horizontally transferred genes (Fisher’s exact tests, P < 0.001). Specifically, genes matching the proMGE database were 7.20-fold enriched, whereas genes on contigs classified as a plasmid or virus were 8.69 and 3.32-fold enriched among HGT genes, respectively. In contrast, genes on contigs classified as containing proviruses were 7.29-fold more likely not to be found among the HGT set.

With perhaps the exception of proviruses, these functional enrichments agree with basic expectations of gene classes expected to be enriched among HGT events in nature.

### Overall relationship between horizontal gene transfer and ecological variables

We next compared how rates of HGT between genomes are associated with co-occurrence and phylogenetic distance (**Supplementary Figure 2**). To do so, we mapped metagenomic reads of previously published ocean metagenomics samples against the 15,339 genomes to determine each genome’s relative abundance across samples. We identified 1,862 samples where at least one genome was called as present. For these samples, a mean of 26,592,809 (SD: 36,857,948) reads were mapped, which represented a mean of 23.9% (SD: 10.2%) of reads per sample. A median of 286 genomes were identified as present across these samples. We restricted our subsequent analyses to 10,388 genomes that were called as present in at least 10 metagenomics samples, and then computed the pairwise co-occurrence of these genomes. This prevalence filtering step meant that 32.3% of genomes were excluded, which resulted in the loss of 54.8% of genomes involved in HGT based on the clustering approach (a drop from 2,295 to 1,037 unique genomes). This indicates that many HGT events occurred in taxa for which we have insufficient data to study their co-occurrence relationships within this dataset (**Supplementary Figure 1**).

As expected, we observed a clear association between taxa co-occurrence and HGT (**Figure 3a**; odds ratio: 19.3; P < 0.001). In contrast, we did not observe the expected negative association between phylogenetic distance and extent of co-occurrence based on the focal genome pairs (**Figure 3b**), and instead there was a near-zero association (Spearman ⍴=0.056; P < 0.001; **Supplementary Figure 3a**). As a reminder, our analysis focused on genome pairs in different genera or above, to more reliably infer HGT events, and these distributions (**Figure 3**) are based on this subset of genome pairs. However, when considering genome pairs within the same genus or lower, which were excluded from our focal analysis, we did find the expected negative association (Spearman ⍴=-0.325; P < 0.001; **Supplementary Figure 3b**). For the subset of genome pairs connected by HGT (**Figure 3b**, right panel) there is lower phylogenetic distance compared to those with no evidence for HGT (mean difference: 0.900;

**Figure 3:**
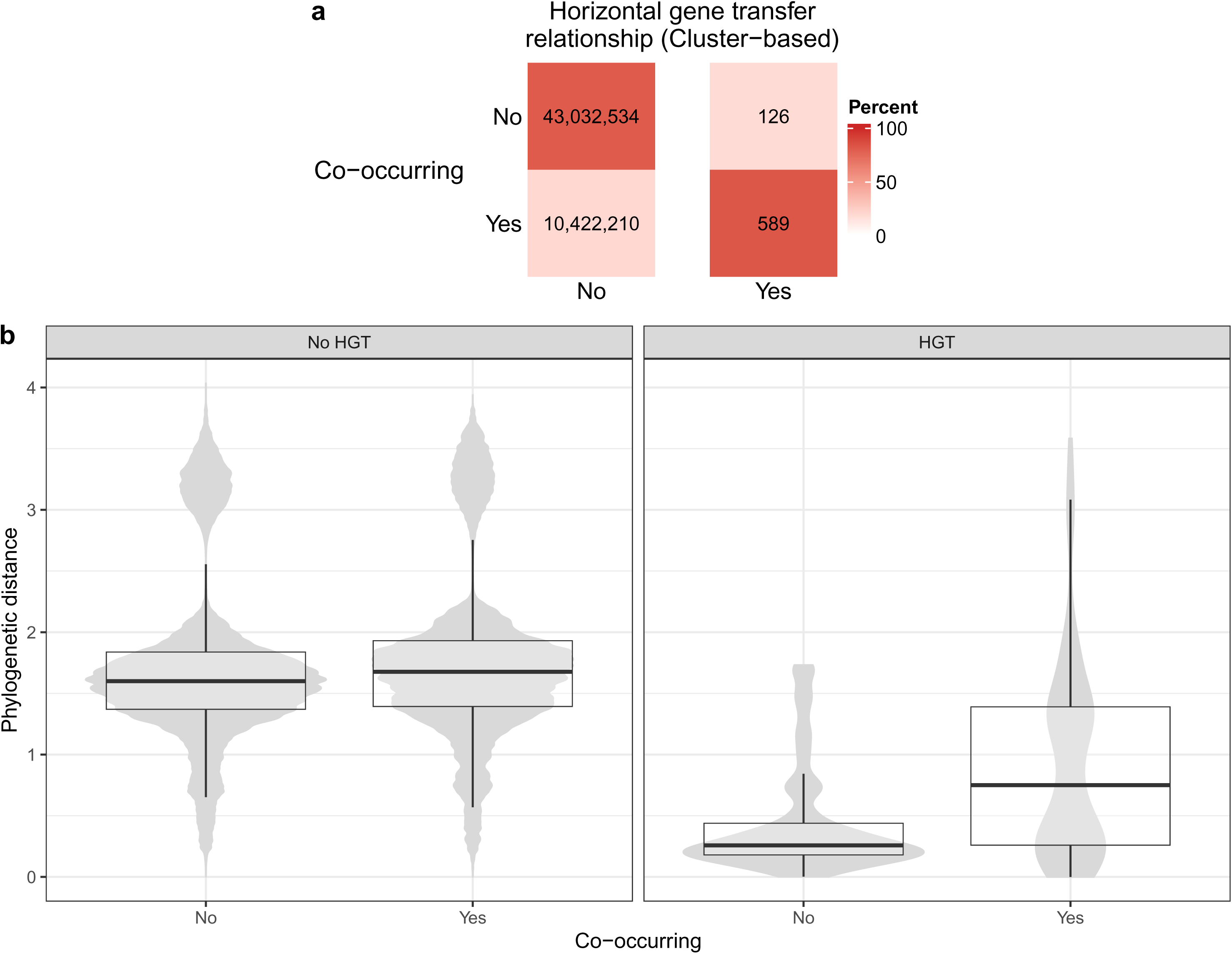
Breakdown of putative horizontally transferred genes identified, by co-occurrence and phylogenetic distance. (a) Tallies and percentages of pairwise genome comparisons, by whether they have at least one horizontal gene transfer (HGT) event inferred and whether they co-occur. (b) Boxplots over violin plots of all genomes tallied in panel a, displaying phylogenetic distance by HGT and co-occurrence. The width of the violin plots does not reflect the total sample size for each category (see panel a for counts in each category).

Wilcoxon test P < 0.001). In addition, genome pairs connected by HGT that do not co-occur have especially low phylogenetic distance compared to those that do (mean difference: 0.485; Wilcoxon test P < 0.001). This last observation could be due to collider bias (i.e., a misleading signal driven by data subdividing; see Discussion).

Overall, we find the expected associations between HGT and both phylogenetic distance and co-occurrence, but the negative association between co-occurrence and phylogenetic distance is only present for closely related taxa.

Phylogenetic distance has previously been proposed as partially explaining the association between co-occurrence and HGT, but other factors may also constrain this link. An HGT event might be adaptive under specific environmental conditions, making it more likely to be retained and observed. Indeed, genome pairs connected by HGT tend to be sampled under more similar environmental conditions compared to genomes without evidence for HGT (**Figure 4**). This result highlights that including information on environmental factors is also needed, in addition to phylogenetic distance, to better interpret the relationship between co-occurrence and HGT.

**Figure 4:**
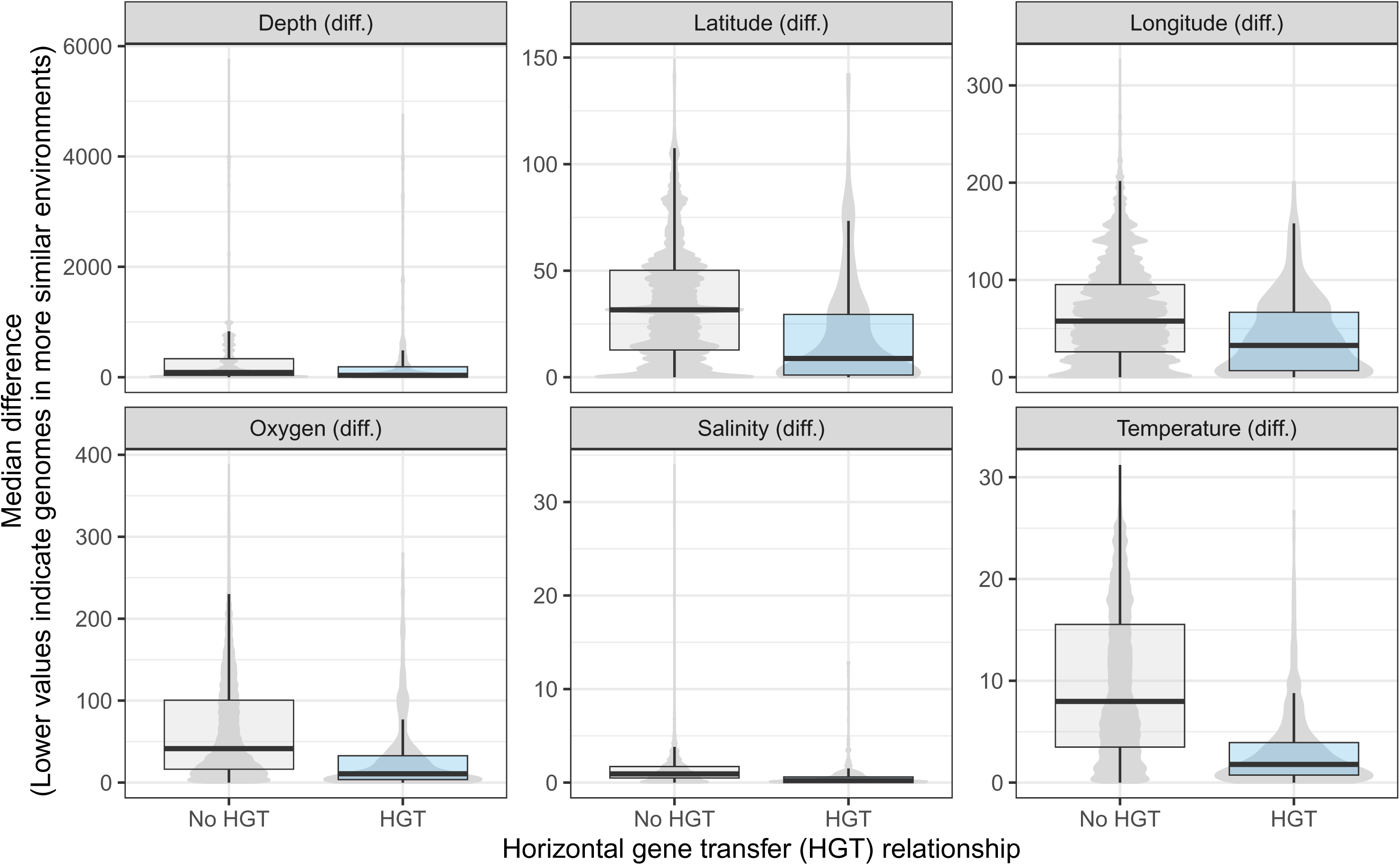
Distributions of the difference in median value of each environmental variable between genome pairs, split by genome pairs connected by HGT and not. The median value for each variable was computed for each genome based on all the metagenomics samples it was found within. The violin plots for the “no HGT” relationship show more detailed variation likely because there are many more genome pairs in this category.

### Testing for a relationship between horizontal gene transfer and co-occurrence, after correcting for phylogeny and environmental factors

We next investigated whether the relationship between HGT and co-occurrence remains, and to what degree, after controlling for phylogenetic distance and median environmental differences. To do so, we built a logistic regression model to predict the binary state of whether the genome pairs are connected by HGT or not. The predictor variables included whether the genome pairs are co-occurring or not, as well as the continuous phylogenetic distances and median environmental differences. Most of the measured environmental factors were abiotic (e.g., temperature, depth), but we also included a biotic size fraction factor as a proxy for genomes associated with free-floating or potentially particle-associated niches, quantified by different filter sizes (see Methods). A lower filter-cutoff enriches for smaller free-floating cells, whereas a larger filter can include particle-attached prokaryotes, which we refer to as “less-filtered” samples (these samples also include free-floating microbes, and are not solely particle-attached). We hypothesized that genomes of particle-associated microbes are more likely to undergo HGT due to higher cell density and potentially higher encounter rates with other taxa.

Based on the model coefficients (**Figure 5**), the clearest predictor of HGT is co-occurrence (2.26, P < 0.001). As this is a binary predictor, this refers to the increased log-odds of being connected by HGT when genomes co-occur, or 9.58-fold (e^2.26^) increased odds. Phylogenetic distance was also highly significant in the model (-1.66, P < 0.001). As this is a continuous variable this means that for every one unit increase in transformed phylogenetic distance (which corresponds to one standard deviation due to the orderNorm transformation), there is a 5.26-fold decrease in HGT. Although the median differences in all environmental variables were also significant, the absolute coefficients are much lower compared to phylogenetic distance. Finally, pairs where both genomes are associated with “less-filtered” metagenomics samples, which are more likely to include particle-attached prokaryotes, were positively associated with HGT (1.26, P=0.013; with wide 95% confidence intervals). Overall, this result is consistent with the relationship between co-occurrence and HGT remaining robust after also including potential ecological drivers.

**Figure 5:**
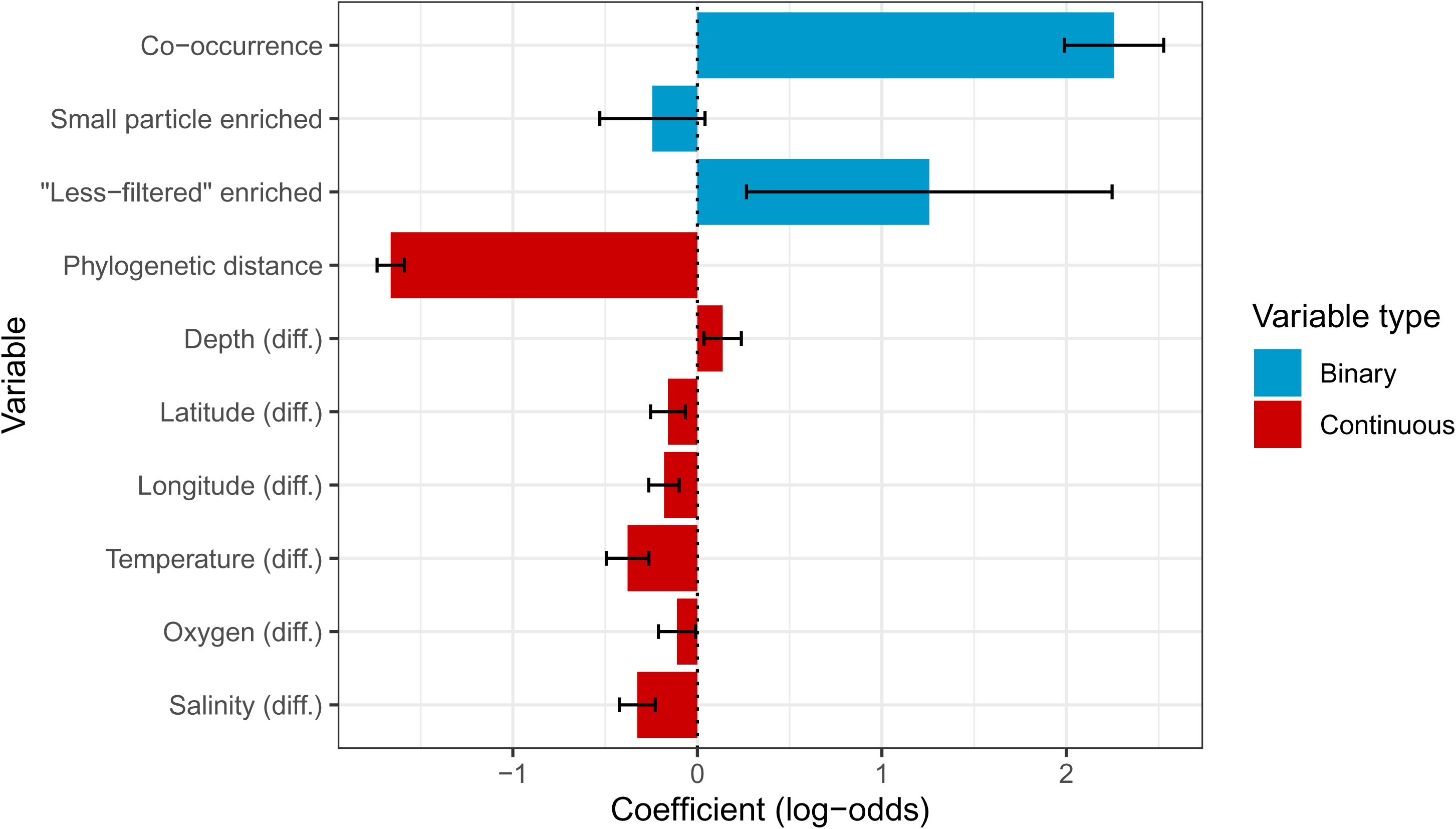
Coefficients in logistic regression model based on all pairwise genome comparisons. The response is whether those genomes have at least one called horizontal gene transfer event (based on the cluster-based approach) between them or not. Co-occurrence (based on the hypergeometric approach) is a binary variable indicating whether those genomes significantly co-occur, and small particle enriched and “less-filtered” enriched indicate whether the genomes being compared are both enriched in samples filtered to enrich for small particles (small), or those with little or no size filters (“less-filtered”). The other variables are ordered quantile normalized continuous values. Variables with ‘diff.’ represent the median inter-sample difference in each environmental variable for all samples encoding the two genomes being compared. Coefficient values less than 0 indicate that decreases in those variables are associated with increases in horizontal gene transfer, whereas a positive value represents the opposite. The intercept for this model was -14.35. Error bars represent 95% confidence intervals (1.96 x standard error). All variables were significant (P < 0.05) except for “small particle enriched”.

We fit this full logistic regression model, and other models based on subsets of predictors. We found the full model fit best based on the Akaike Information Criterion (AIC; difference in AIC of -3.46 compared to the next best model). In addition, although there is potential for multicollinearity between these variables, the variance inflation factor was < 2 in all cases, meaning that this is not a concern for this model.

The above analyses focused on results from our focal bioinformatics workflow, but we also ensured our overall results did not change if differing methods for determining co-occurrence or HGT were used, or if different datasets were analyzed. Accordingly, we re-ran this logistic regression analysis based on two other methods for determining co-occurrence and two other methods for determining HGT, as well as with genomes from the proGenomes database (**Supplementary Figure 4**; see Methods). Although the differing HGT-identification methods have similar motivations, they can result in highly different profiles. For instance, identifying HGT with RANGER-DTL between strains of a species resulted in a huge number of putative events identified (the positive intercept for these models indicating that more than half of genome comparisons were called to have at least one HGT, or recombination, event). Nonetheless, in almost all cases, the models which included all tested variables produced the best-fitting models.

Exceptions included the model based on cluster-based HGT and ‘simple’ co-occurrence, as well as all models based on the proGenomes database, where the models excluding the particle size groupings fit best. In the case of the proGenomes database models, this is likely due to few genomes found to be associated with each grouping for this genome database (see Methods).

Overall, this broader investigation resulted in the same major interpretations: phylogenetic distance and co-occurrence are strongly associated with HGT, even after including additional factors. One exception to our general interpretation is a slight negative association (coefficient of -0.13) between co-occurrence and HGT in the model based on BLASTn-identified HGT events and propr-based co-abundance (**Supplementary Figure 4**; see Discussion). Nonetheless, we also found overall concordance of our results based on an independent genome database (proGenomes), from which we excluded metagenomics-derived assemblies, and corresponded to higher quality genomes than our focal set (although with fewer inferred HGT events).

Finally, we also ran additional checks to help ensure our interpretations are not driven by technical artifacts. We re-ran several models on the largest subsets of the overall OceanDNA metagenomics dataset: samples from *Tara* Oceans and GEOTRACES separately (**Supplementary Figure 5**). Our goal here was to ensure that dataset substructure was not driving our results. The model coefficients were qualitatively similar compared to the models on the entire dataset (with one exception, but which included few HGT events and had high multicollinearity). We also ran generalized linear mixed models to control for the genome IDs in each comparison (i.e., to control for individual genomes which could be disproportionately driving the observed signals). To do so, we duplicated the input table used for focal models, to allow each genome ID to be associated with each comparison separately. Due to this approach, the error estimates for these models are less reliable compared to the original logistic regressions. Nonetheless, these models provide insight into how controlling for genome ID can affect the models, and we found that our overall interpretations remained consistent (**Supplementary Figure 5**). The exception was that the association trends for the filter-cutoff groupings associated with genome pairs were less consistent (see Discussion). This is not unexpected, as there were only four genome pairs associated with “less-filtered” samples and called for HGT in the original analysis. These pairs consisted of only seven unique genomes (**Supplementary Table 1**).

### In-depth environmental associations with horizontal gene transfer

Our main analysis focused on which factors are associated with HGT occurrence between pairs of genomes. This was useful for investigating how co-occurrence and phylogenetic distance relate to HGT, but the relationship to environmental factors was less clear. We thus conducted follow-up analyses, focused on samples rather than genome pairs, to investigate the association with more specific environmental factors. We did so by computing a per-sample measure of the prevalence of HGT: the proportion of genomes in each sample with at least one putative HGT event identified.

To characterize the extent of association between broader oceanographic environmental factors and HGT prevalence per sample, we acquired detailed environmental data for a subset of our *Tara* Oceans samples (see Methods). We focused on 12 environmental factors across 131 metagenomic samples (all small particle size samples; see Methods). We first characterized the variation in these variables across the samples, and identified clear clustering of environmental variables, such as depth with photosynthetically available radiation (**Supplementary Figure 6**). We then built a random forest model to predict HGT prevalence based on these variables (R^2^ = 0.687). Based on a permutation approach we assessed the significance of variable importance (**Figure 6a**), and identified chlorophyll *a*, photosynthetically available radiation (PAR), bbp470, nitrate, oxygen, fluorescent Coloured Dissolved Organic Matter (fCDOM), and depth all as significant predictors. This was expected as these variables show clear associations with horizontal gene transfer prevalence individually (**Supplementary Figure 7**). In particular, there is a discrete clustering of samples by low and high PAR, which are associated with higher and lower HGT prevalence, respectively (**Figure 6b**).

**Figure 6:**
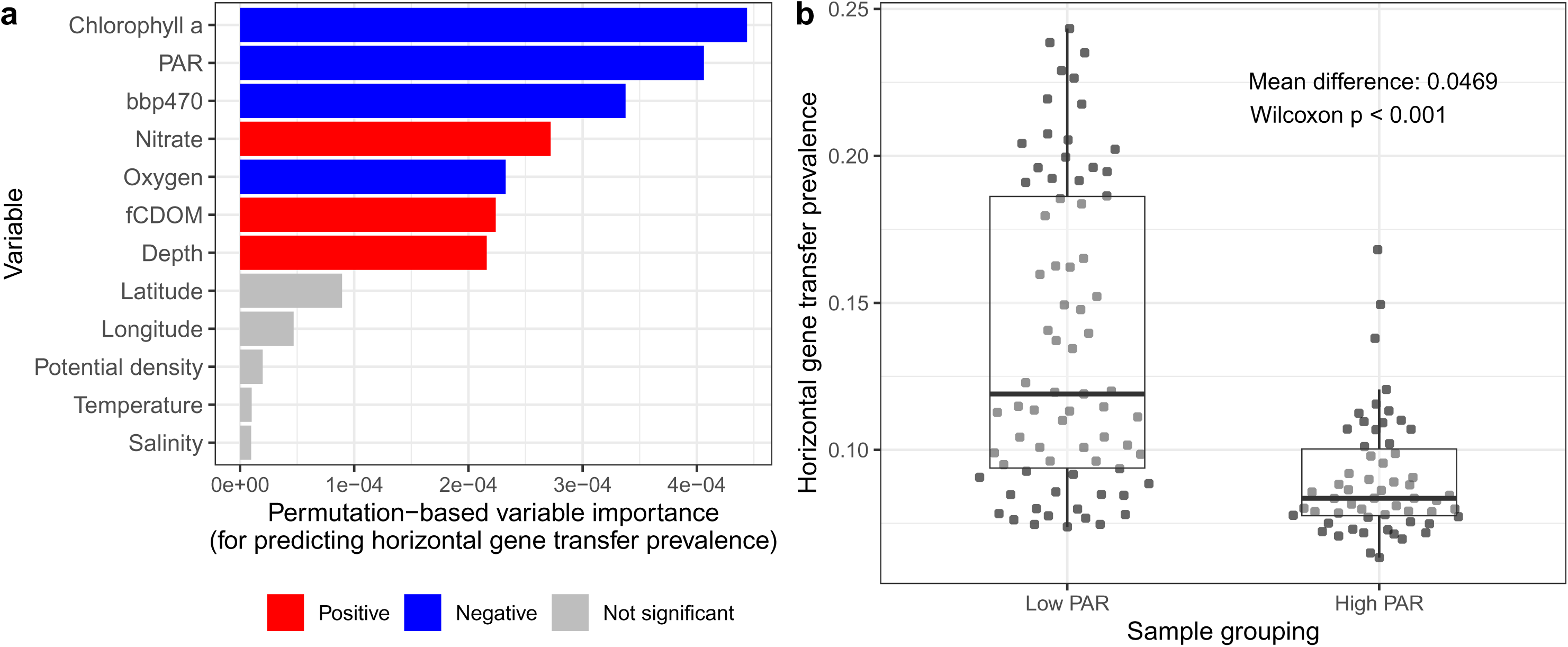
Key associations between environmental variables and horizontal gene transfer prevalence. (a) Variable importance of tested environmental variables for predicting horizontal gene transfer prevalence. Variable colour indicates the direction of the association (based on Spearman correlation, for context), while grey indicates variables that did not have significant variable importance. (b) Boxplots of horizontal gene transfer prevalence for samples split into different groupings of PAR (lower and above 20%, which is the approximate centre of a large range with no samples).

For improved insight into each variable’s effect in the model, we also investigated the Individual Conditional Expectation plots (**Supplementary Figure 8a**). These highlight that each variable has only a small impact on the predicted HGT prevalence (i.e., there is a small range in how much the predicted HGT prevalence varies when each environmental variable is systematically altered). In addition, there are clear signals of the predictive value for chlorophyll *a*, PAR, and bbp470 mainly being driven by the difference between values near 0 (with high predicted HGT prevalence) and larger values for these variables (with low predicted HGT prevalence). Overall, there is a consistent effect on all samples’ predictions as the variables are altered, indicating that these variables primarily have independent impacts on predicted HGT prevalence. Nonetheless, there are interactions between variables that impact HGT prevalence (**Supplementary Figure 8b**), particularly among oxygen, fCDOM, and nitrate, for which pairwise interactions explained on average 10.6% of their joint effects.

### Paired comparison of small and large particle enriched samples

Our models focusing on genome pairs found a weak signal that genomes in ‘less-filtered’ samples undergo more HGT. However, such samples also include free-floating bacteria and are confounded by dataset divisions (e.g., almost all ‘less-filtered’ samples included were from the GEOTRACES dataset). Accordingly, we conducted a similar follow-up on whether HGT prevalence, and related features, show clear differences across samples processed with differing size fractions. We did so by focusing on additional *Tara* Oceans metagenomics samples that matched a subset of those analyzed above, but which were restricted to a higher size fraction (with small particles excluded, unlike the earlier ‘less-filtered’ grouping; see Methods). Since these are the same collected samples, this allowed us to evaluate the effect of filter size fraction while implicitly controlling for environmental factors (**Figure 7**). We found a clear signal of higher HGT prevalence in higher size fraction samples (a 2.09-fold increase on average), with similar signals for genomes with hits to the proMGE database or to genes matched to plasmids or viruses, but no significant difference for genomes containing proviruses.

**Figure 7:**
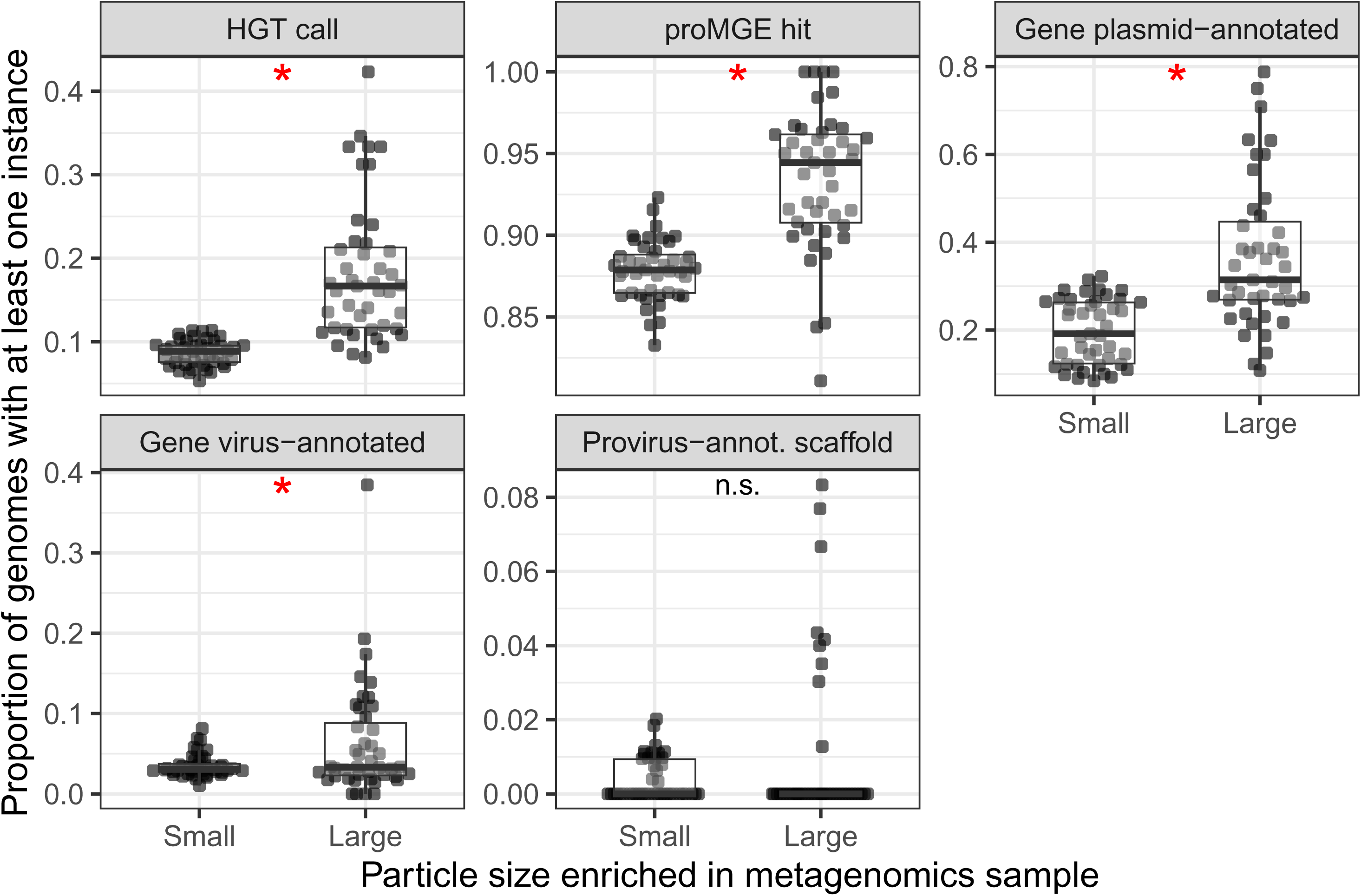
Breakdown of horizontal gene transfer (HGT) and related feature prevalence across 43 *Tara* Oceans samples. The same biological samples are matched based on whether they were enriched for small or large particles based on the fraction filter (i.e., to enrich for free-floating or particle-attached bacteria). The y-axis for all panels is the proportion of genomes in a sample with at least one instance of that category (e.g., the proportion of genomes in a sample with at least one gene identified to be either acquired or transferred through HGT). Asterisks indicate significant comparisons (adjusted P < 0.05) based on paired Wilcoxon tests.

## Discussion

We have shown that co-occurring environmental genomes in ocean metagenomics data are more likely to show evidence for HGT. In agreement with an earlier study using lower-resolution 16S rRNA gene amplicon sequencing data [11], we found that the association between co-occurrence and HGT holds after controlling for phylogenetic distance and several measured environmental variables. This result underlines the strong association between co-occurrence and HGT.

Our work also agrees with a survey of antibiotic resistance genes in metagenomes across a wide range of environments, which similarly identified decreased HGT with phylogenetic and environmental distance [84]. In this recent study, the authors identified not only phylogenetic distance as negatively associated with HGT, but specifically genetic incompatibility, which was based on several factors, including the difference in 5-mer content between genomes. Intriguingly, cases were identified where genomes with low genetic incompatibility displayed high rates of HGT despite being phylogenetically distant. Although we did not specifically investigate genetic incompatibility here, this will be an important area for future investigation to determine whether similar trends are found to explain general HGT patterns, and not when restricted to antibiotic resistance genes.

Through our main workflow for identifying HGT events, we were able to stratify genes putatively shared through HGT by taxonomic level and sequence identity (as a proxy for time since transfer). The functional enrichment of these genes largely agreed with expectations from the literature (**Figure 2**). However, several functional enrichments were in the opposite direction from previously reported expectations. Particularly for enrichments based on genes at >= 99% identity, we found several functional enrichments in opposite directions at different taxonomic levels. For instance, amino acid transport and metabolism was depleted for transfer events at the genus level, but enriched at the phylum and domain levels. These taxonomic level-specific enrichments would be missed if functional enrichment were performed on all genes combined, and could reflect differing recurrent selective pressures for HGT at varying taxonomic levels.

The association between HGT and both co-occurrence and phylogenetic distance is extremely clear in our data (**Figure 1** and **Figure 3**). However, the expected association between co-occurrence and phylogenetic distance was only present in genome pairs within the same genus or lower. This observation highlights that closely related taxa are primarily driving this signal, rather than a consistent trend at all phylogenetic distances, at least in this marine dataset. Additionally, we identified an intriguing trend where genome pairs that co-occur and show evidence for HGT were significantly more phylogenetically distant compared to those that do not co-occur yet undergo HGT (**Figure 3b**). This could indicate more complex associations between co-occurrence, HGT, and phylogenetic distance than previously appreciated. For instance, this observation could be driven by metabolic complementation across genomes, which is known to be associated with co-occurrence [85], and could be enriched for distantly related genomes in our dataset. However, a technical explanation for this observation, which we think is more likely, is collider bias [86]. When both phylogenetic distance and co-occurrence independently affect the probability of genomes being connected by HGT, conditioning on one factor (e.g., co-occurrence) creates an expected difference in the distribution of the other factor (e.g., phylogenetic distance) among genome pairs. In other words, if genome pairs can be connected by HGT either due to being phylogenetically similar or due to co-occurring, then we would expect that genome pairs connected by HGT that are not co-occurring would have lower phylogenetic distance on average.

There are at least two biological mechanisms that could underlie the general observation of an association between HGT and co-occurrence. First, taxa that are physically closer could simply have more opportunities for HGT, which must be true to some extent. For instance, taxa in completely isolated environments, with no gene flow or possibility of DNA transfer, cannot undergo HGT. In regions of the ocean stratified by geography, depth, and environmental factors, it is reasonable to expect that taxa would undergo HGT more frequently with other taxa sharing the same local environment.

Second, an alternative causal explanation is that HGT of habitat-specific genes allows species sharing those genes to thrive in that habitat, and thus to co-occur. In this alternative explanation, co-occurrence is a proxy for unmeasured environmental variables that define cryptic microbial habitats. Distinguishing between these explanations for the observed associations is an avenue for future research, potentially harnessing time-series sampling to infer causal relationships.

Although we cannot formally distinguish between these explanations, our results are consistent with both physical proximity driving HGT and, potentially, HGT contributing to shared niche preferences. We consistently found evidence for increased HGT among genomes in larger particle-associated filter fractions. This suggests that physical proximity on particles, in contrast to reduced encounter rates among planktonic bacteria enriched on the small filter fractions, could drive HGT. The larger filter fraction could also share common selective pressures, associated with a particle-attached lifestyle, which may also explain the association with HGT independently of physical proximity. Consistent with environmental drivers of HGT (or that HGT of niche-adaptive genes allows survival in similar environments), we found that similarity in a few measured environmental factors was associated with higher rates of HGT in logistic regression models. However, the effects of these environmental factors on HGT was relatively small compared to the larger effects of co-occurrence and phylogeny.

Based on a more in-depth analysis for a small subset of samples with additional environmental data we identified clear environmental factors associated with HGT. Disentangling the causal factors underlying these associations, such as for chlorophyll *a*, which is a marker of phytoplankton biomass, generally higher at depths with optimum nutrient and light conditions (e.g., deep chlorophyll maxima), is difficult.

However, we can infer that low-light, nitrate-rich environments are enriched for HGT. This work agrees with the previously identified association between increasing depth (or at least variables associated with increased depth) and transposase genes [87, 88]. In both our analyses and a previous study [81], transposase genes were also associated with genomes sampled from the large particle enriched filter fraction. This highlights the challenges of disentangling the effects of physical proximity and environmental similarity on HGT rates. It may also explain why co-occurrence – which effectively captures both the effects of proximity and niche similarity in both measured and unmeasured niche dimensions – is so strongly associated with HGT.

A novel aspect of our study is that we focused on metagenomic sequencing data, in a complementary fashion to the approach of analyzing 16S ribosomal RNA gene sequencing profiles and matching these to reference genomes [11]. However, most genomes in our focal dataset were MAGs, which have contamination and misassembly issues [89, 90], including chimeric MAGs which might not be detected based on standard quality controls. In addition, depending on species clonality within a given metagenomic sample, MAGs may represent a population genome for a species, which may bias and preclude the detection of HGT across MAGs. To address these problems, we conducted an additional analysis based on high quality genomes from the proGenomes database, with MAGs excluded. Our overall interpretations were similar (**Supplementary Figures 4** and 5), suggesting that our inferences of HGT were not driven by misassembled MAGs.

We also considered a variety of bioinformatic workflows for detecting HGT and determining co-occurrence. One clear observation was that within-species inferences of HGT and recombination with RANGER-DTL led to an extremely high number of putative hits. Because phylogenetic reconciliation approaches are sensitive to errors in tree-building [91], we cannot reject that this observation is simply driven by inaccuracies in the gene tree topologies. Indeed, we are skeptical that these hits are reliable inferences for exact HGT events, but nonetheless these results produced similar model outputs as our other approaches. In one model we found results that disagreed with the overall interpretation that co-occurrence and HGT are positively associated: a model based on BLASTn-inferred HGT events and propr co-abundance (**Supplementary Figure 4**). We focused on the cluster-based HGT events, as BLASTn-inferred events reflect a network of potential HGT partners. Specifically, all genomes that share genes above a certain identity cut-off were considered HGT connected based on the BLASTn results, even if a better match exists. In addition, unlike co-occurrence based on presence/absence of genomes, the propr results are based on genome relative abundances. There could be a bias where similar genes are difficult to independently assemble into separate MAGs when these genes are present in genomes at similar abundances in metagenomics data. In any case, all other models, including those based on more straightforward data, agreed with our general interpretation.

In conclusion, we have shown that the relationship between co-occurrence and HGT is robust to phylogeny and broad environmental factors. However, we also observed increased HGT between genomes sampled under more similar environmental conditions, and among particle-associated genomes. Future work disentangling the importance of physical proximity (at microscopic or larger scales) compared to more specific shared environmental pressures in driving HGT in the global ocean is needed.

## Acknowledgements

We would like to thank all members of the Bobay, Chaffron, and Shapiro labs for feedback.

