## Supplemental Materials for "Co-occurrence is associated with horizontal gene transfer across marine bacteria independent of phylogeny"

### Supplementary Materials

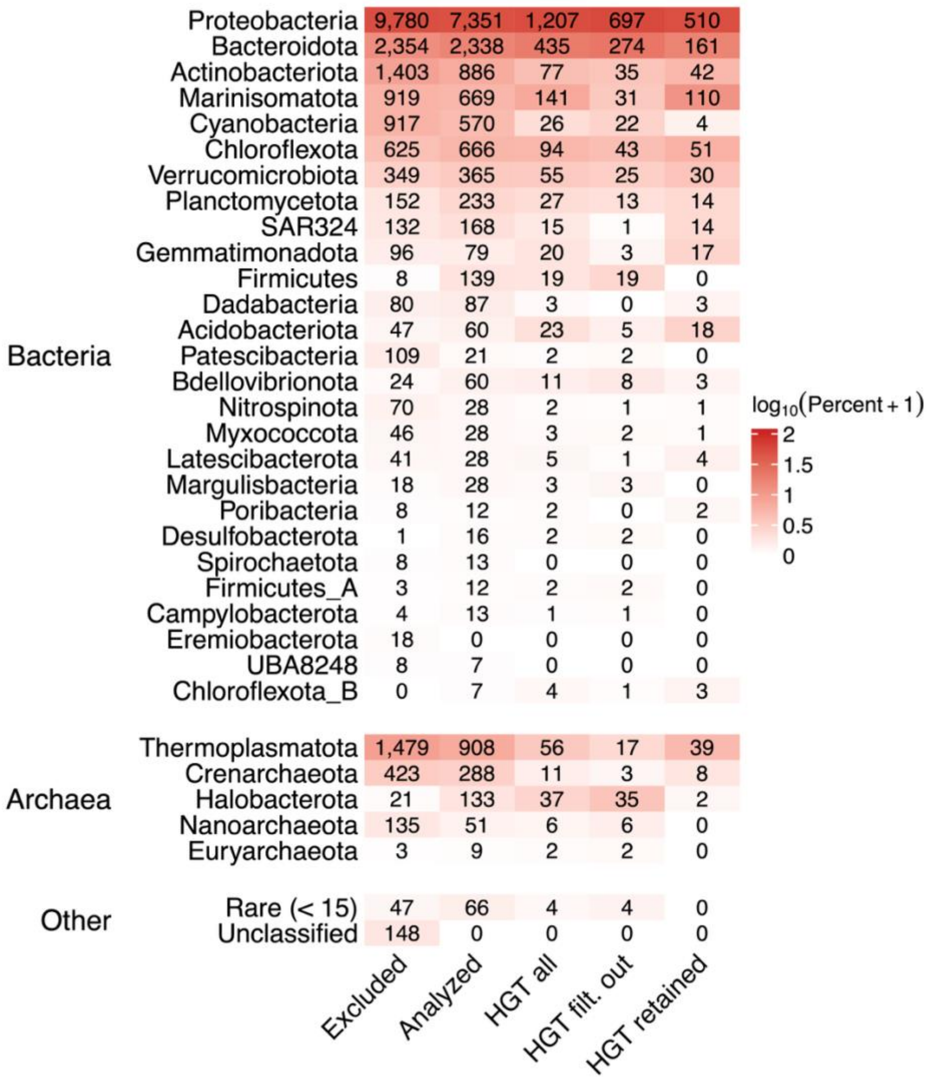


**Supplementary Figure 1**: Counts of analyzed genomes by phylum, split into the bacterial and archaeal domains, and coloured by percentage. Only genomes from the focal genome database are shown. “Excluded”: genomes excluded due to being low quality. “Analyzed”: the set of all genomes analyzed in this study. “HGT all”: all genomes with at least one putative horizontal gene transfer (HGT) event identified based on the clustering approach (which are a subset of “Analyzed”). The “HGT filt. out” and “HGT retained” groupings represent genomes that were subsets of “HGT all”, and were either below or above the metagenomics sample prevalence cut-off, respectively. In other words, they represent genomes that were lost or retained, respectively, after requiring genomes to be present in 10 samples based on mapping metagenomics reads to genomes with CoverM.

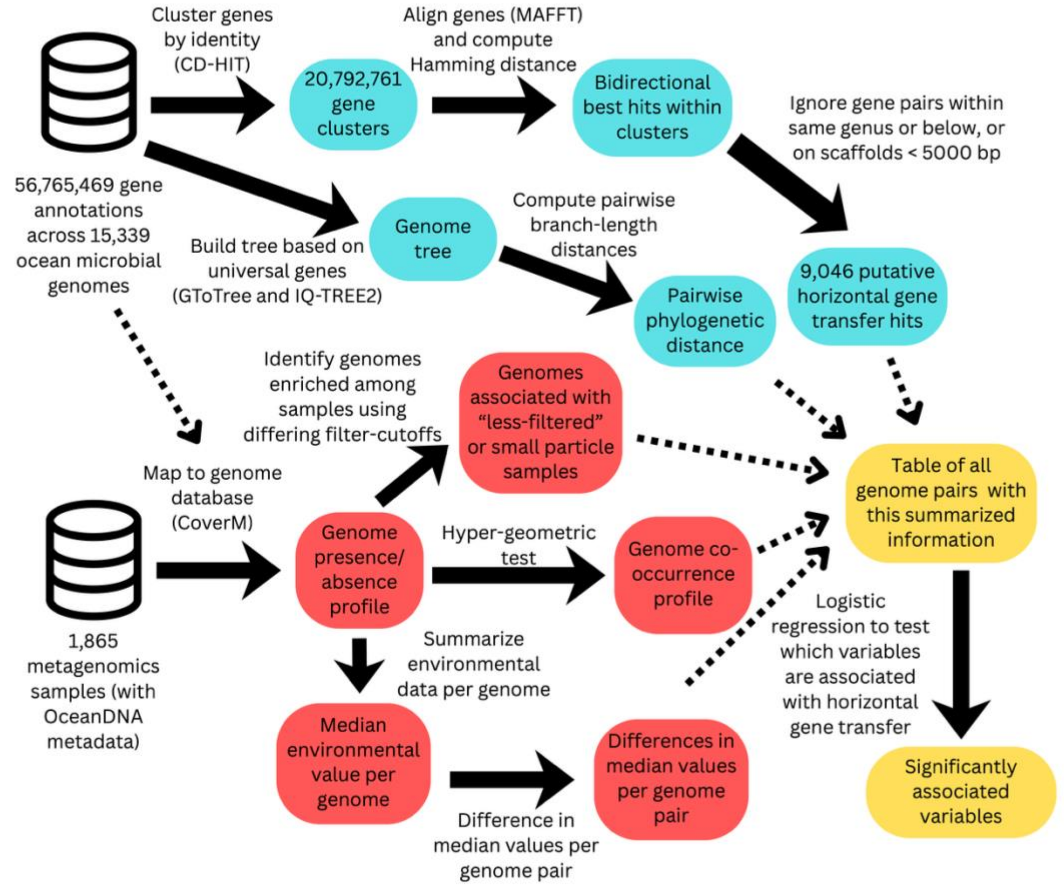


**Supplementary Figure 2**: Focal analysis workflow used in this study, which involved identifying putative horizontal gene transfer events based on clustering genes and identifying co-occurring genomes with a hyper-geometric test. Supplementary workflows were based on this structure, but with different steps for identifying horizontal gene transfer events and co-occurring genomes.

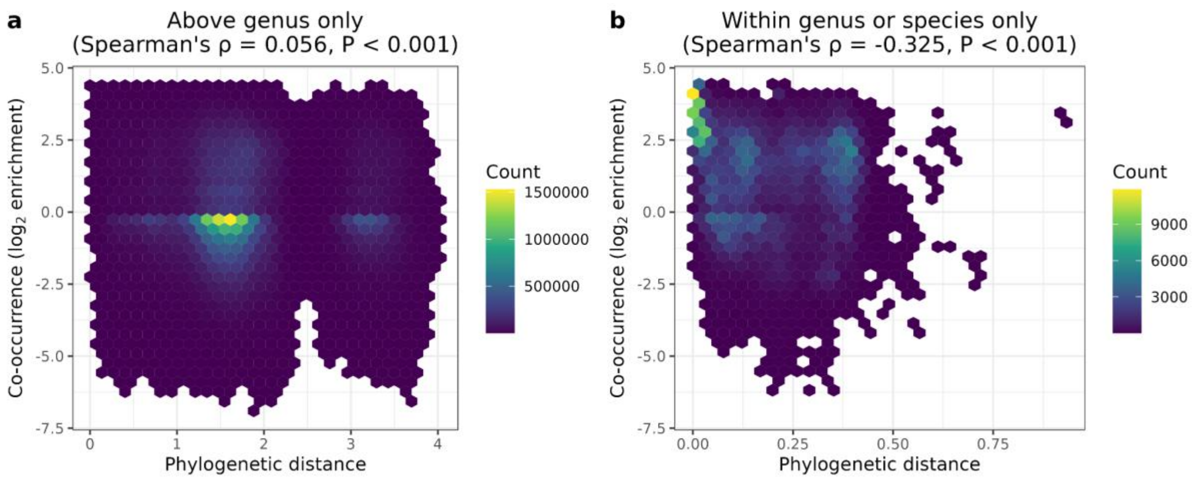


**Supplementary Figure 3**: Hex plots (coarse scatterplots with coloured hexes indicating counts of overlapping points) of the co-occurrence and phylogenetic distance for all pairwise genome comparisons in the focal dataset. (a) Restricted to genome comparisons between different genera and higher (which we focused on). (b) Restricted to genome comparisons between genomes in the same species or genus (which we excluded from all other analyses for our focal workflow). The plotted co-occurrence metric is: $\log_2((\text{Observed} + 1)/(\text{Expected} + 1))$, where "Observed" is the number of samples observed to share both genomes based on CoverM mapping, and "Expected" is the number expected by chance, based on the prevalence of each genome. Spearman correlation results are indicated above each panel. For the Spearman correlation tests, simply the ratio of Observed/Expected was used as the co-occurrence metric.

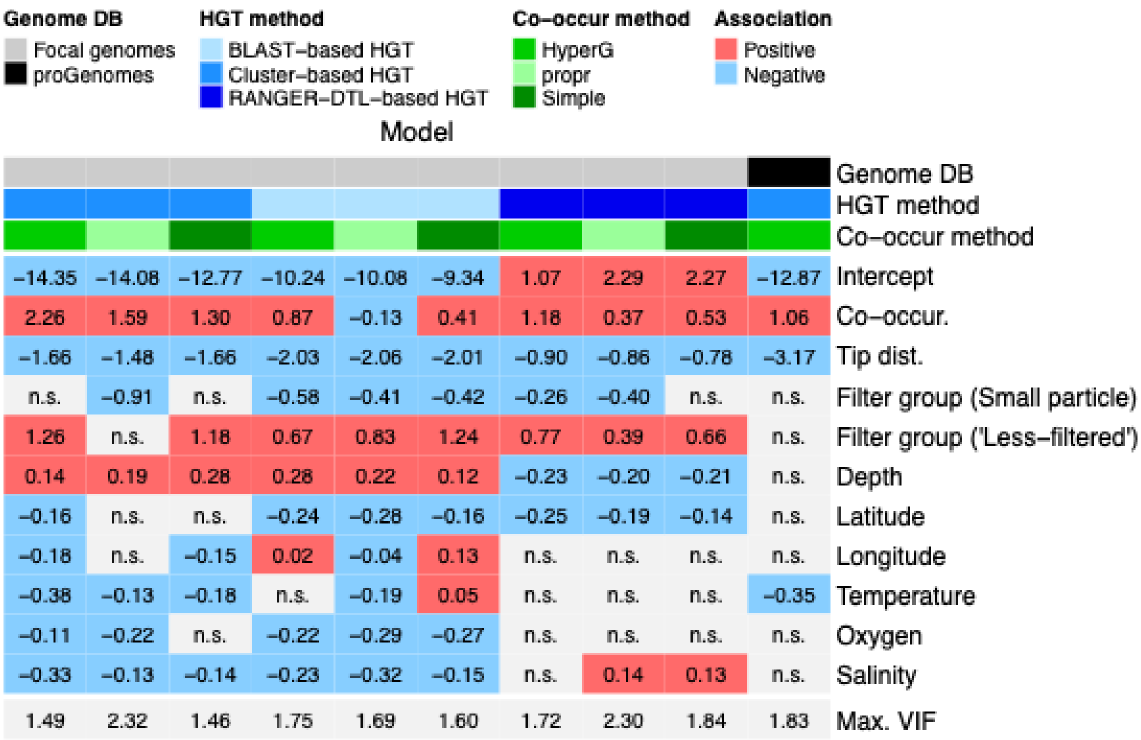


**Supplementary Figure 4**: Coefficients in logistic regression models based on all pairwise genome comparisons, as in Figure 5 but for a wide range of input data. The first column corresponds to the focal model, displayed in Figure 5. Each column is a different model, with input data described by the top three coloured strips: the genome database (DB), horizontal gene transfer (HGT)-detection method, and co-occurrence method. All coloured cells in the heatmap represent significant coefficients per model, and their direction. Non-significant coefficients (P >= 0.05) are indicated by "n.s.". The final row is the maximum variance inflation factor (Max. VIF) per model, indicating that the input variables are moderately, but not highly, correlated. The variable types (binary vs. continuous) are the same as Figure 5, which affects the interpretation of these coefficients, except for co-occurrence in the simple and propr-based models, which was continuous. In addition, all environmental variables tested refer to the median difference in environmental variables.

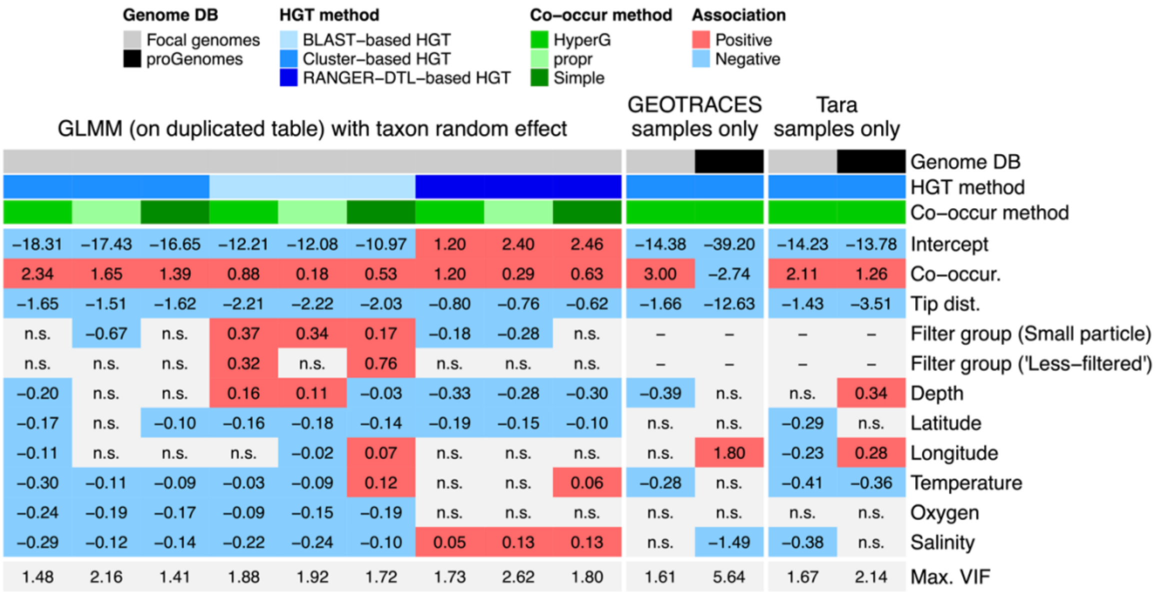


**Supplementary Figure 5**: Similar to Supplementary Figure 4, but for additional model outputs. The first nine columns are results for generalized linear mixed models (GLMMs) with genome ID as the random effect. The last four columns are model coefficients for logistic regression models based on the subset of datasets with only GEOTRACES or *Tara* Oceans samples. Non-significant coefficients (P >= 0.05) are indicated by “n.s.”. The final row is the maximum variance inflation factor per model (Max. VIF), which indicates that most models do not have concerning levels of multicollinearity, except for the proGenomes GEOTRACES-only model with a value > 5, indicating high multicollinearity. There was also an extremely low intercept for this model (-39.20), which reflects there being few HGT events identified for genomes in this subset.

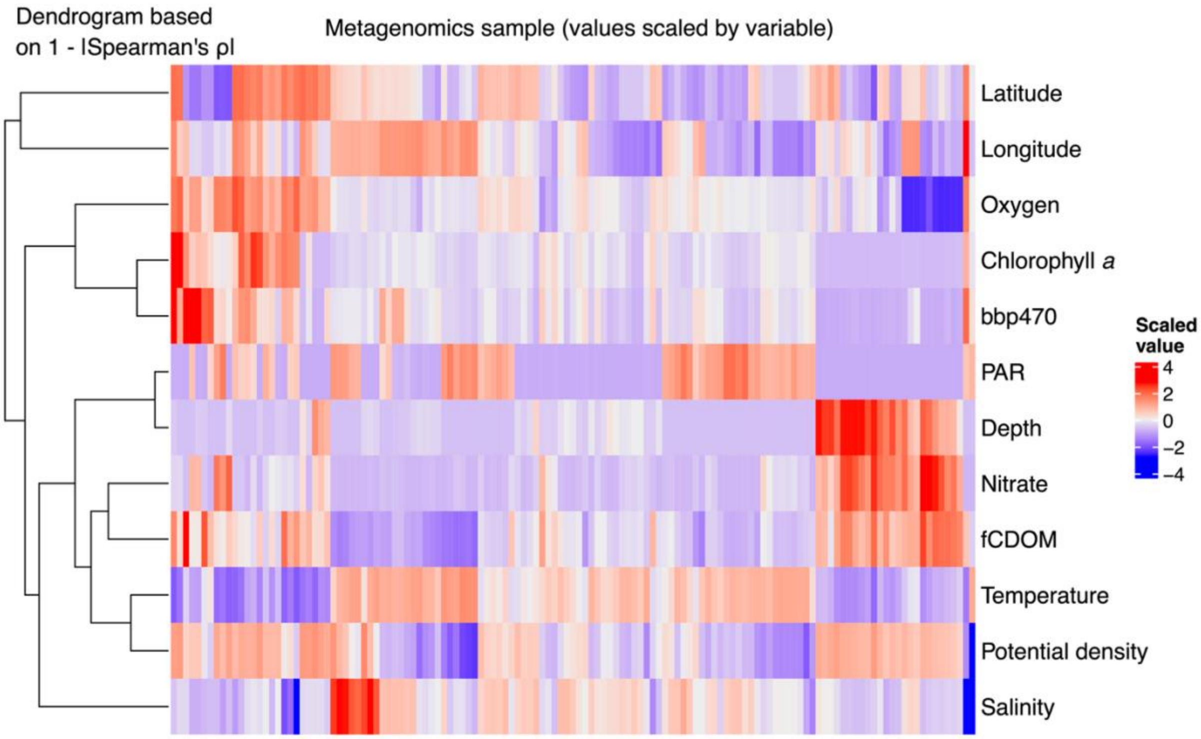


**Supplementary Figure 6**: Heatmap of environmental variables (rows) across 131 *Tara* Oceans samples (columns). Variables were clustered based on the complement of their pairwise absolute Spearman correlation coefficients (ρ), while samples were clustered based on pairwise Euclidean distances. Hierarchical clustering was performed using the average linkage method. All environmental variables were mean centred and scaled by the standard deviation prior to this analysis.

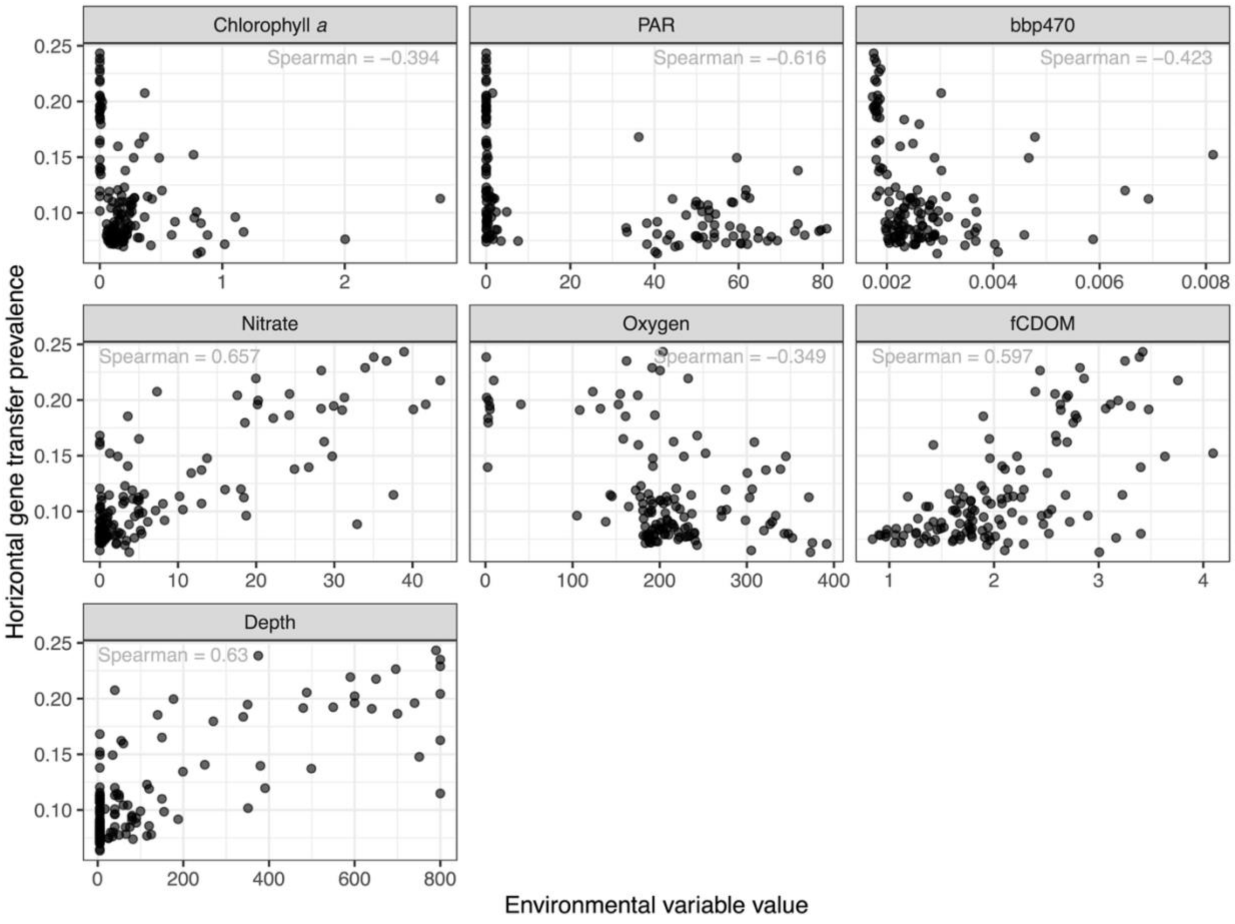


**Supplementary Figure 7**: Scatterplots between environmental variables and HGT prevalence. Restricted to factors significant in random forest model only. The Spearman correlation coefficient is indicated on each panel (all P < 0.001). Variables are ordered by their variable importance in the random forest model (from highest to lowest).

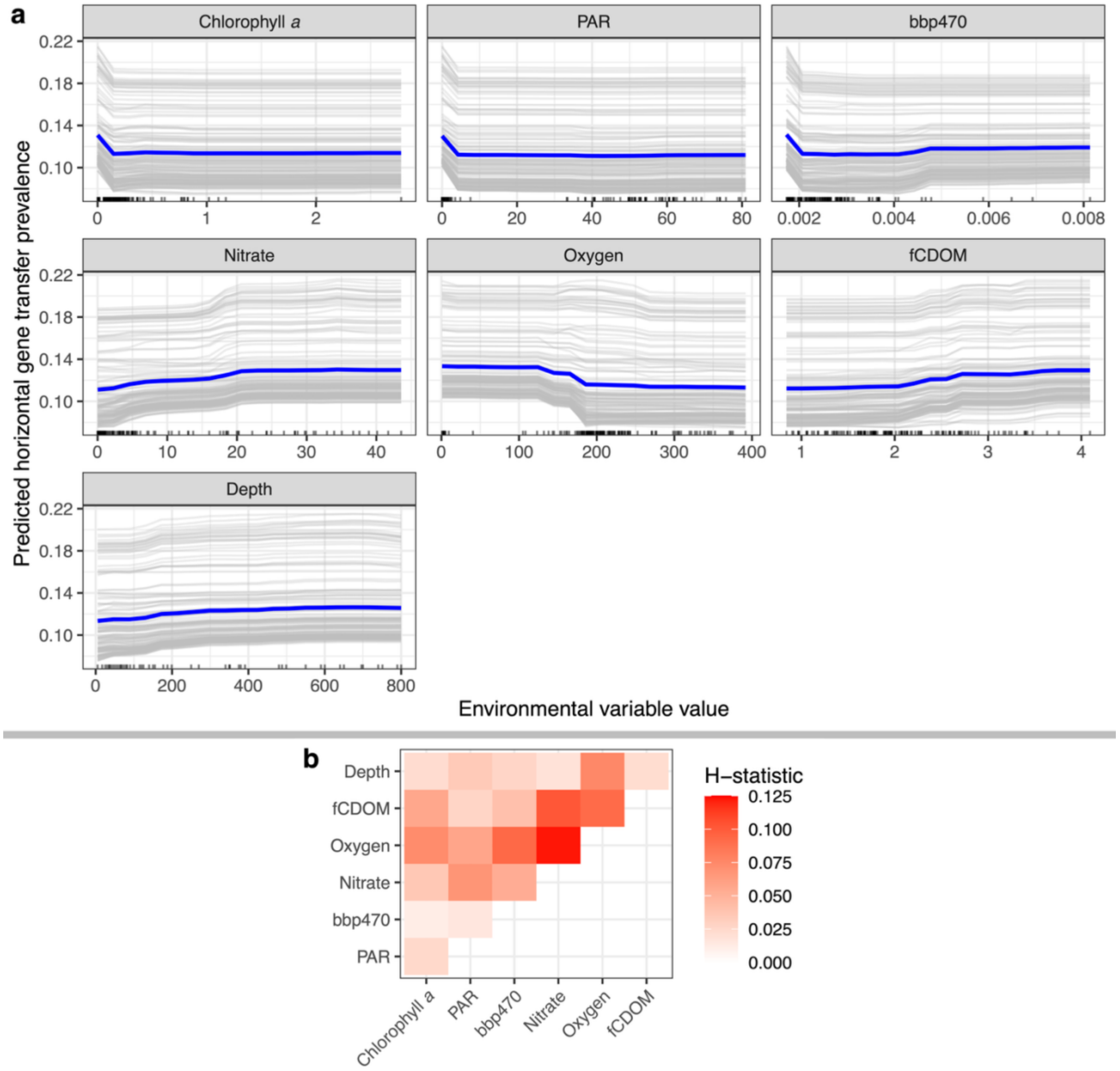


**Supplementary Figure 8**: (a) Individual Conditional Expectation plots for environmental variables that significantly predict HGT prevalence in random forest models. These plots show how predictions (for horizontal gene transfer prevalence, on the y-axis) change across varying values of each predictor. For each sample (grey lines) all other environmental variables are held at their observed values while the focal environmental variable is systematically altered (across x-axis). Blue lines indicate the average of all samples and black ticks at the bottom indicate the observed samples. The exact thresholds for how these lines shift are not reliable in regions with few samples. For instance, the exact threshold where increased photosynthetically active radiation (PAR) results in lower predicted values (the initial dip) is not reliable as there is a large PAR range with no samples (centred on 20). We can only infer that lower PAR values are associated with higher predictions, but more sampling is needed to identify exact thresholds. (b) Pairwise interactions between significant variables in the random forest model, represented by the H-statistic. This statistic corresponds to the proportion of the joint effect of each pair of variables that is explained by their interaction (e.g., about 12% of the joint effect of oxygen and nitrate is explained by their interaction).

**Supplementary Table 1**: Taxonomy of seven metagenome-assembled genomes driving signal of positive association between genomes enriched in 'less-filtered' metagenomics samples and horizontal gene transfer

| **Study-assigned taxon ID** | **Taxonomy** |
|---|---|
| Taxa_11628 | d__Bacteria; p__Chloroflexota; c__Dehalococcoidia; o__UBA3495; f__UBA3495; g__UBA9611; s__; TARA_SAMEA2623756_METAG_DIFAIHGK |
| Taxa_3418 | d__Bacteria; p__Actinobacteriota; c__Acidimicrobiia; o__Microtrichales; f__MedAcidi-G1; g__UBA9410; s__; MALA_SAMN05422166_METAG_OAKDMEED |
| Taxa_8602 | d__Bacteria; p__Bacteroidota; c__Bacteroidia; o__Flavobacteriales; f__Flavobacteriaceae; g__MED-G13; s__; TARA_SAMEA2621990_METAG_OMFNCAAD |
| Taxa_442 | d__Bacteria; p__Chloroflexota; c__Dehalococcoidia; o__UBA3495; f__UBA3495; g__UBA3495; s__UBA3495 sp002716645; BGEO_SAMN07136539_METAG_DOCKDDBM |
| Taxa_86 | d__Bacteria; p__Chloroflexota; c__Dehalococcoidia; o__UBA3495; f__UBA3495; g__UBA3495; s__UBA3495 sp002716645; BATS_SAMN07137087_METAG_CDOPEKMJ |
| Taxa_892 | d__Bacteria; p__Gemmatimonadota; c__Gemmatimonadetes; o__SG8-23; f__UBA6960; g__UBA1138; s__UBA1138 sp003447875; BGEO_SAMN07136727_METAG_BJDLPBMF |
| Taxa_8654 | d__Bacteria; p__Bacteroidota; c__Bacteroidia; o__Flavobacteriales; f__Flavobacteriaceae; g__MED-G14; s__; TARA_SAMEA2622074_METAG_COJFLNPO |
